# Deep Brain Stimulation rescues the homeostasis disruption of circulating D- and L-amino acids level in men with Parkinson’s Disease

**DOI:** 10.1101/2025.08.13.668904

**Authors:** Tommaso Nuzzo, Federica Carrillo, Marcello Serra, Claudia Gentile, Anna Di Maio, Alessandra Pizzella, Sara Pietracupa, Nicola Modugno, Francesco Errico, Teresa Esposito, Alessandro Usiello

**Author notes:** contributed equally and share joint first authorship (Tommaso Nuzzo, Federica Carrillo, Marcello Serra). to whom correspondence should be addressed: Corresponding authors: Dr Teresa Esposito PhD Molecular Genetics and Genomics Laboratory Institute of Genetics and Biophysics, “Adriano Buzzati Traverso”, Italian National Research Council (CNR) 80131 Naples – Italy. PI IGB-CNR laboratory IRCCS INM Neuromed 8077 Pozzilli – Italy., Prof Alessandro Usiello PhD. Department of Environmental, Biological and Pharmaceutical Sciences and Technologies (DISTABIF). Università della Campania, L. Vanvitelli, Viale Abramo Lincoln 5, 81100, Caserta, Italy. PI Neuroscience Laboratory. CEINGE, Biotecnologie Avanzate Franco Salvatore, S.c.a.rl. 80145 Naples-Italy.

## Abstract

Recent evidence indicates a marked downregulation of circulating D- and L-amino acids involved in regulating glutamatergic NMDAR function in Parkinson’s disease (PD) patients compared with matched controls. However, the extent to which disease progression and antiparkinsonian therapies contribute to this dysregulation remains unclear.

To address these issues, in the present study we measured by High Performance Liquid Chromatography the concentrations of glutamatergic system-related D- and L-amino acids and their precursors in the plasma of male and female healthy controls (HC) and PD patients across three distinct clinical stages and treatment conditions: (1) early stage L-DOPA naïve patients treated with MAO-B inhibitors; (2) mid-stage patients treated with L-DOPA; and (3) advanced stage patients receiving Deep Brain Stimulation in the subthalamic nucleus (STN-DBS) plus L-DOPA.

Our results reveal notable reduction of circulating neuroactive D- and L-amino acids exclusively in male PD patients, while female patients exhibit a similar directional trend. In male patients, this dysregulation manifests early, with L-DOPA–naïve individuals showing decreased plasma levels of L-glutamate and L-aspartate. In mid-stage L-DOPA-treated PD patients, amino acid reductions extend to L-alanine, L-serine, L-glutamine, L-asparagine, and L–threonine. Remarkably, in advanced PD patients, with a median disease duration of ∼ 23 years, STN–DBS normalizes the blood concentrations of these amino acids to those observed in HC.

In conclusion, our study highlights the potential of circulating D- and L-amino acid dysregulation as an early biomarker of PD and demonstrates that, in contrast to L-DOPA therapy, the STN-DBS confers systemic metabolic benefits even at advanced stages of the disease.

## Introduction

Parkinson’s disease (PD) is a progressive neurodegenerative disorder characterized by α-synuclein accumulation and the loss of dopaminergic neurons in the substantia nigra pars compacta (SNpc) (Bloem et al., 2021).

PD is clinically characterized by cardinal motor symptoms—bradykinesia, rigidity, and resting tremor— alongside a range of non-motor symptoms such as sleep disturbances, anosmia, and mood alterations, with cognitive impairment typically emerging in later stages (Obeso et al., 2017). Although the majority of PD cases are sporadic, approximately 5–10% have a defined genetic aetiology (Jankovic and Tan, 2020).

Currently, there is no cure for PD, and standard treatment primarily relies on administration of the dopamine precursor L-DOPA. Although L-DOPA is effective in alleviating motor symptoms in the short term, its long-term use is associated with several adverse effects, including dyskinesias, motor response fluctuations, and neuropsychiatric complications (De Deurwaerdère et al., 2017).

These limitations have driven the search for alternative strategies, with deep brain stimulation (DBS) of the subthalamic nucleus (STN) or globus pallidus internus emerging as a prominent option. DBS has demonstrated substantial efficacy in alleviating parkinsonian motor symptoms and reducing L-DOPA-associated motor side effects in individuals with advanced PD (Anderson et al., 2017). However, its effect on non-motor symptoms remains limited and heterogeneous (Anderson et al., 2017).

Glutamatergic transmission alterations, particularly those involving NMDA receptors (NMDAR), are known to be involved in PD neuropathology (Campanelli et al., 2022). Previous studies have shown that modulation of NMDAR activity via the NR1 allosteric site—using endogenous amino acids such as glycine (Gly) and D-serine (D-Ser) or the partial agonist D-cycloserine—can produce therapeutic effects on motor, non-motor, and cognitive symptoms in both rodent and primate models of PD, as well as in patients (Frouni et al., 2021; Gelfin et al., 2012; Ho et al., 2011; Pawlak et al., 2012; Schmitz et al., 2013).

Through Ultra-Performance Liquid Chromatography-Mass Spectrometry (UPLC-MS) and High-Performance Liquid Chromatography (HPLC) analyses, we recently demonstrated significantly increased levels of the NMDAR co-agonist D-Ser, and its precursor L-serine (L-Ser), in the putamen of 1-methyl-4-phenyl-1,2,3,6-tetrahydropyridine (MPTP)-intoxicated monkeys, in the *post-mortem* caudate-putamen of PD patients, and in the cerebrospinal fluid (CSF) of *de novo* PD patients compared to controls (Di Maio et al., 2023; Gervasoni et al., 2025a).

Further supporting dysregulation of amino acids involved in glutamatergic transmission and energy metabolism in PD, our recent metabolomic analysis of serum from 69 consecutive PD patients and 32 healthy controls—combining untargeted Proton Nuclear Magnetic Resonance (^1^H-NMR) and targeted UPLC/MS— revealed elevated glutamine and serine plus reduced α-ketoglutarate concentrations in PD patients compared to controls by NMR, alongside a pronounced reduction in glutamic acid levels by UPLC/MS (Gervasoni et al., 2025b). Of great relevance, we also demonstrate that these serum alterations are significantly modulated by sex and genetic background (Yahyavi et al., 2025). Indeed, in a well-characterized cohort of 121 idiopathic PD patients, 124 carriers of pathogenic LRRK2, PARK2, PINK1, PARK7, TMEM175, or GBA1 PD-related mutations, and 203 healthy controls, our HPLC analysis revealed a remarkable male-specific reduction in circulating NMDAR-related amino acids and their precursors, particularly in idiopathic cases (Yahyavi et al., 2025).

While these findings reinforce the concept of PD as a multisystem disorder (Costa et al., 2023; Liu et al., 2025), it remains unclear whether the observed serum metabolic alterations reflect basal ganglia pathology or stem from peripheral organ dysfunction. Moreover, it has yet to be determined whether and how these changes are affected by the disease stage and treatment, since all PD cases in our previous studies had a median disease duration of 5–6 years and were receiving L-DOPA (Gervasoni et al., 2025b; Yahyavi et al., 2025).

To address these critical issues, in the present study we measure by HPLC the concentration of glutamatergic system-related D- and L-amino acids and their precursors in the plasma of healthy controls (HC) and PD patients across three distinct clinical stages and treatment conditions: 1) L-DOPA-naïve (n = 16; median disease duration 1.5 years), L-DOPA-treated (n = 15 median disease duration 7 years), and STN-DBS-treated (n = 14; median disease duration 23 years). Our study identifies the dysregulation of several neuroactive amino acids as an early potential metabolic fingerprint for PD and demonstrates that STN-DBS confers systemic amino acids metabolism benefits in its advanced stages.

## RESULTS

### Clinical features of the study cohort

16 HC and 45 PD patients were considered in this study. Demographic and clinical features of sex-stratified participants are reported in **Table 1**. PD individuals were categorized into three subgroups based on sex, pharmacological treatment and clinical characteristics: L-DOPA-naïve (male N = 11, female N = 5; undergoing MAO-B inhibitor treatment), L-DOPA (male N = 11, female N = 4; undergoing L-DOPA treatment), and DBS (male N = 9, female N = 5; undergoing L-DOPA treatment with STN-DBS) (**Table 1**).

**Table 1.**
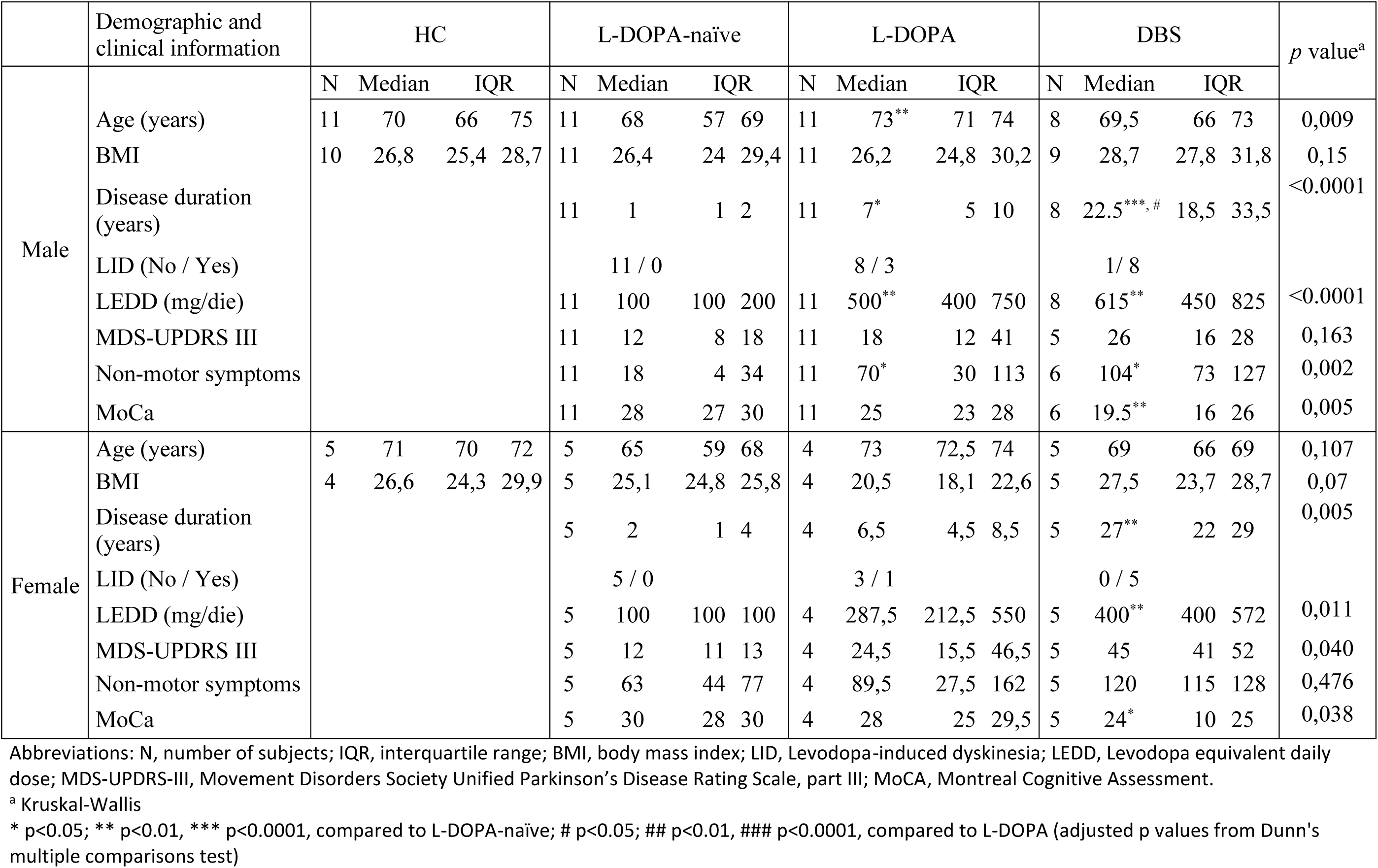
Demographic and clinical parameters of Parkinson’s diesease and heathy control subjects enrolled in the study.

The age of PD patients and HC was comparable across groups irrespective of sex, except for males in L-DOPA group, who were significantly older than those without L-DOPA treatment (**Table 1**). BMI did not differ significantly across groups; however, a trend toward higher values was observed in DBS-treated patients of both sexes (**Table 1**). Male and female DBS-treated patients showed a longer duration of the disease, irrespective of sex (**Table 1 and Fig. 1**). Both L-DOPA- and DBS-treated groups of either sex were characterized by higher levodopa equivalent daily dose (LEDD) and presence of patients manifesting L-DOPA-induced dyskinesia (LID) (**Table 1 and Fig. 1**). Regarding clinical features, no significant differences in motor symptoms (assessed by UPDRS Part III) were observed across male PD patient groups (**Table 1 and Fig. 1**). However, male patients in the L-DOPA and DBS groups exhibited a greater burden of non-motor symptoms (NMS), and those in the DBS group showed increased cognitive impairment (as measured by MoCA), compared to sex-matched L-DOPA–naïve patients (**Table 1 and Fig. 1**). In female patients, motor and NMS did not differ significantly among treatment groups; however, female patients receiving DBS displayed greater cognitive impairment relative to sex-matched L-DOPA–naïve patients (**Table 1 and Fig. 1**).

**Figure 1.**
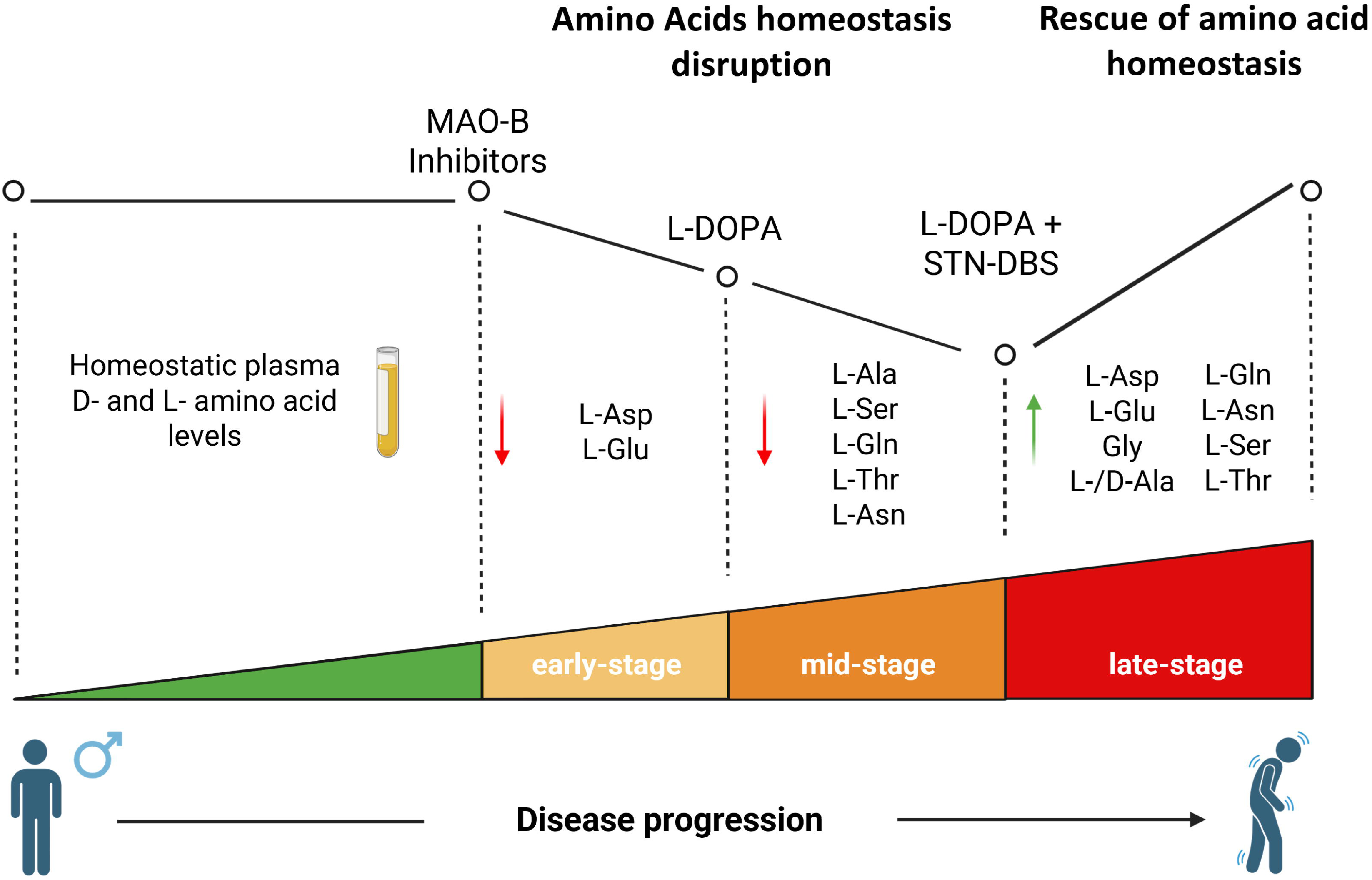
Clinical profiles of male and female Parkinson’s disease patients treated with MAO-B inhibitors, L-DOPA, or deep brain stimulation plus L-DOPA. **(a, b, f, g)** Comparison of disease duration (a, f) and LEDD (**b, g**) among PD patients without L-DOPA treatment (L-DOPA-naïve, male, N=11; female, N= 5), PD patients with L-DOPA therapy (male, N=11; female, N= 4), or PD patients with deep brain stimulation (DBS; male, N=8; female, N= 5) in male (**a, b**) and female (**f, g**) groups. (**c-e; h-j**) MDS-UPDRS III, non-motor symptoms and MOCA scores among PD patients without L-DOPA treatment (L-DOPA-naïve; male, N=11; female, N= 5), PD patients with L-DOPA therapy (L-DOPA; male, N=11; female, N= 4), or PD patients with deep brain stimulation (DBS; male: MDS-UPDRS N=5; NMS and MOCA, N=6; female, N= 5) in male (**c-e**) and female (**h-j**) groups. Data are shown as violin plots representing the median with interquartile range. Dots represent values from each patient analyzed. *p < 0.05, **p<0.01, ***p <0.0001, compared to L-DOPA-naïve; # p < 0.05, compared to L-DOPA (Dunn’s post hoc multiple comparisons).

### STN-DBS therapy rescues circulating NMDAR-related amino acid levels in men with PD

We first examined by HPLC how PD condition, including treatment and disease stage, influenced plasma levels of NMDAR-related D- and L-amino acids—including L-glutamate (L-Glu), L-aspartate (L-Asp), glycine (Gly), D-alanine (D-Ala), and D-serine (D-Ser)—in male and female HC and PD groups.

In male PD subjects, non-parametric Kruskal–Wallis analyses revealed significant alterations of plasma L-Glu, L-Asp, Gly, D-Ala, and D-Ser concentrations (**Fig. 2**; **Table 2**). These findings were further confirmed by ANCOVA on natural log-transformed data, adjusted for age and LEDD, to account for potential confounding factors (**Table 2**).

**Figure 2.**
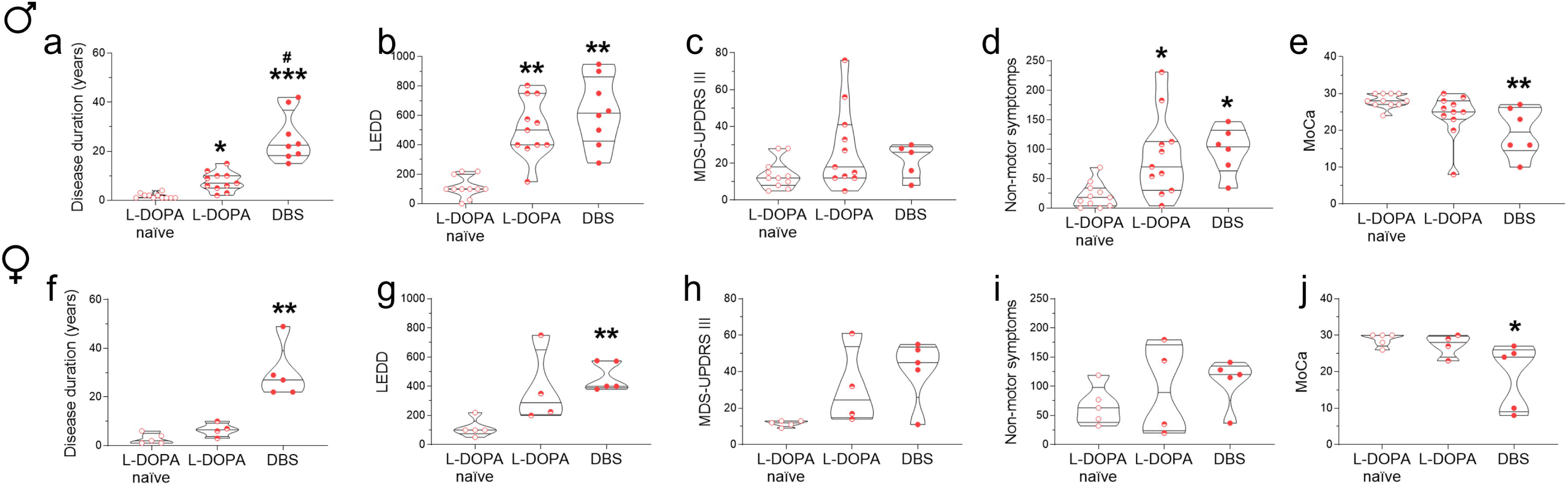
Effect of DBS treatment on the plasma concentrations of NMDAR-related D- and L-amino acids of male and female PD patients. **(a)** Representative HPLC chromatogram obtained from plasma of PD patients with D-serine and D-alanine peaks magnification. **(b-k)** Levels of L-glutamate (L-Glu) (**b, g**), L-aspartate (L-Asp) (**c, h**), glycine (Gly) (**d, i**), D-alanine (D-Ala) (**e, j**) and D-serine (D-Ser) (**f, k**) in the plasma of male (**b-f**) and female (**g-k**) healthy controls subjects (HC; male N=11, female N=5), PD patients without L-DOPA treatment (L-DOPA-naïve; male N=11, female N=5), PD patients with L-DOPA therapy (L-DOPA; male N=11, female N=4), or PD patients with deep brain stimulation (DBS; male N=9, female N=5). Data are shown as violin plots representing the median with interquartile range. Dots represent values from each patient analyzed. *p < 0.05, **p<0.01 (Dunn’s post hoc multiple comparisons).

**Table 2.**
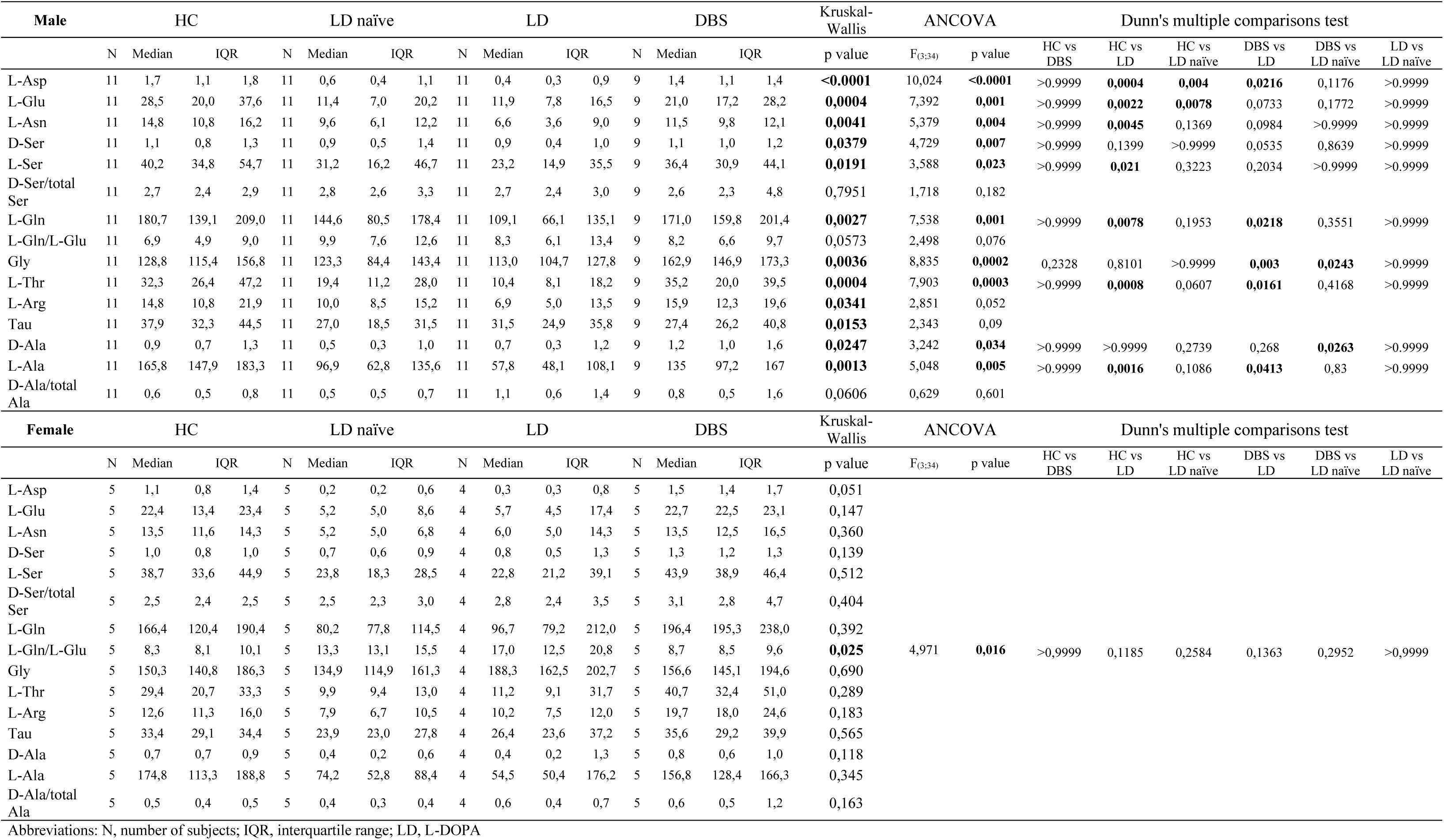
Concentrations of amino acids compared among PD and HC subjects stratified in male and female groups.

Specifically, Dunn’s multiple comparisons test revealed a significant reduction in plasma L-Glu and L-Asp levels in PD patients, both with and without L-DOPA treatment, compared to HC (**Fig. 2**; **Table 2**). A similar trend toward reduced D-Ser levels was observed in L-DOPA–treated PD patients relative to HC, although this difference did not reach statistical significance (p=0.139; **Fig. 2**; **Table 2**).

Notably, male DBS-treated PD patients exhibited higher plasma concentrations of L-Glu, L-Asp, Gly, D-Ala, and D-Ser compared to both L-DOPA–naïve and L-DOPA–treated patients, with levels approaching those observed in HC (**Fig. 2**; **Table 2**). Specifically, Dunn’s multiple comparisons test revealed a significant elevation in plasma D-Ala levels in DBS-treated patients compared to L-DOPA–naïve patients, in L-Asp levels compared to L-DOPA–treated patients, and in Gly levels compared to both L-DOPA–naïve and L-DOPA– treated patients (**Fig. 2**; **Table 2**). Furthermore, in male DBS-treated patients, we observed a trend toward increased plasma L-Glu levels compared to both L-DOPA–naïve (p = 0.177) and L-DOPA–treated (p = 0.073) patients, as well as higher D-Ser levels compared to L-DOPA–treated patients (p = 0.053) (**Fig. 2**; **Table 2**).

Similarly to male patients, female PD patients exhibited reductions in plasma NMDAR-related D- and L-amino acids in both L-DOPA–naïve and L-DOPA–treated groups compared to HC, with levels normalizing following DBS. However, unlike males, these changes did not reach statistical significance in females. Specifically, Kruskal–Wallis tests revealed only trend-level effects for L-Glu (p = 0.147) and L-Asp (p = 0.051) (**Fig. 2**; **Table 2**).

### STN-DBS therapy restores the plasma levels of L-Gln, L-Asn, L-Ala, and L-Thr in men with PD

Given the prominent effects of PD condition on circulating levels of amino acids involved in NMDAR signaling, we next investigated whether similar alterations also extend to other neuroactive D- and L-amino acids not directly associated with NMDAR activation, including L-glutamine (L-Gln), L-asparagine (L-Asn), L-serine (L-Ser), L-alanine (L-Ala), and L-threonine (L-Thr).

In male PD subjects, we reported significant alterations of plasma L-Gln, L-Asn, L-Ser, L-Ala, and L-Thr concentrations (**Fig. 3**; **Table 2**). These findings were further confirmed by ANCOVA on natural log-transformed data, adjusted for age and LEDD (**Table 2**).

**Figure 3.**
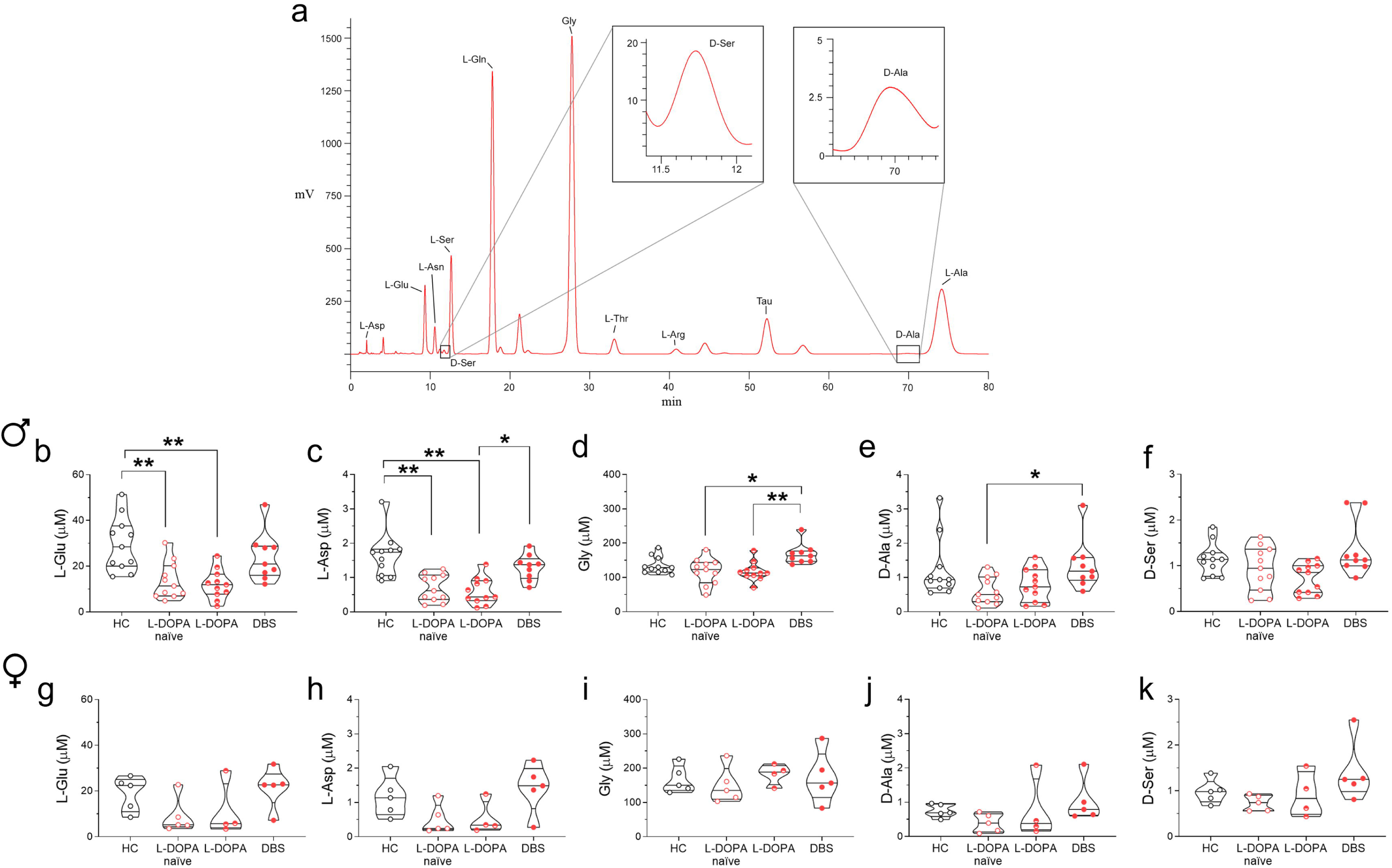
Effect of DBS treatment on the plasma L-glutamine, L-asparagine, L-serine, L-alanine, and L-threonine concentrations in male and female PD patients. Levels of L-glutamine (L-Gln) (**a, f**), L-asparagine (L-Asn) (**b, g**), L-serine (L-Ser) (**c, h**), L-alanine (L-Ala) (**d, i**), and L-threonine (L-Thr) (**e, j**) in the plasma of male (**a-e**) and female (**f-j**) healthy controls subjects (HC; male N=11, female N=5), PD patients without L-DOPA treatment (L-DOPA-naïve; male N=11, female N=5), PD patients with L-DOPA therapy (L-DOPA; male N=11, female N=4), or PD patients with deep brain stimulation (DBS; male N=9, female N=5). Data are shown as violin plots representing the median with interquartile range. Dots represent values from each patient analyzed. *p < 0.05, **p<0.01 (Dunn’s post hoc multiple comparisons).

Notably, Dunn’s multiple comparisons test revealed a significant reduction in the plasma levels of all tested neuroactive amino acids in L-DOPA–treated patients compared to HC (**Fig. 3**; **Table 2**). In contrast, L-DOPA– naïve patients exhibited only non-significant trends toward reduced levels of L-Asn (p = 0.136), L-Ala (p = 0.108), and L-Thr (p = 0.060) relative to HC (**Fig. 3**; **Table 2**). Consistent with previous findings, male DBS-treated PD patients exhibited higher plasma concentrations of all tested amino acids compared to both L-DOPA–naïve and L-DOPA–treated patients, with significantly elevated levels of L-Gln, L-Ala, and L-Thr compared to L-DOPA–treated patients (**Fig. 3**; **Table 2**). Additionally, a non-significant trend toward increased L-Asn levels was observed in DBS-treated patients compared to L-DOPA–treated patients (p = 0.098; **Fig. 3**; **Table 2**).

In contrast to male patients, female PD subjects showed no statistically significant alterations in plasma levels of L-Gln, L-Asn, L-Ser, L-Ala, and L-Thr, regardless of treatment or disease stage. Although L-DOPA–naïve and L-DOPA–treated women demonstrated reductions relative to sex-matched HC—and levels normalized following DBS—these changes did not reach significance by Kruskal–Wallis analysis (Fig. 3; Table 2).

Overall, our findings indicate that the homeostasis of neuroactive D- and L-amino acids is disrupted early— and exclusively—in male PD patients receiving MAO-B inhibitors, with further deterioration observed as the disease progresses under L-DOPA therapy (**Fig. 4**). Notably, in advanced PD patients, STN-DBS combined with L-DOPA effectively restores amino acid profiles to levels comparable to those of healthy controls (**Fig. 4**).

**Figure 4.**
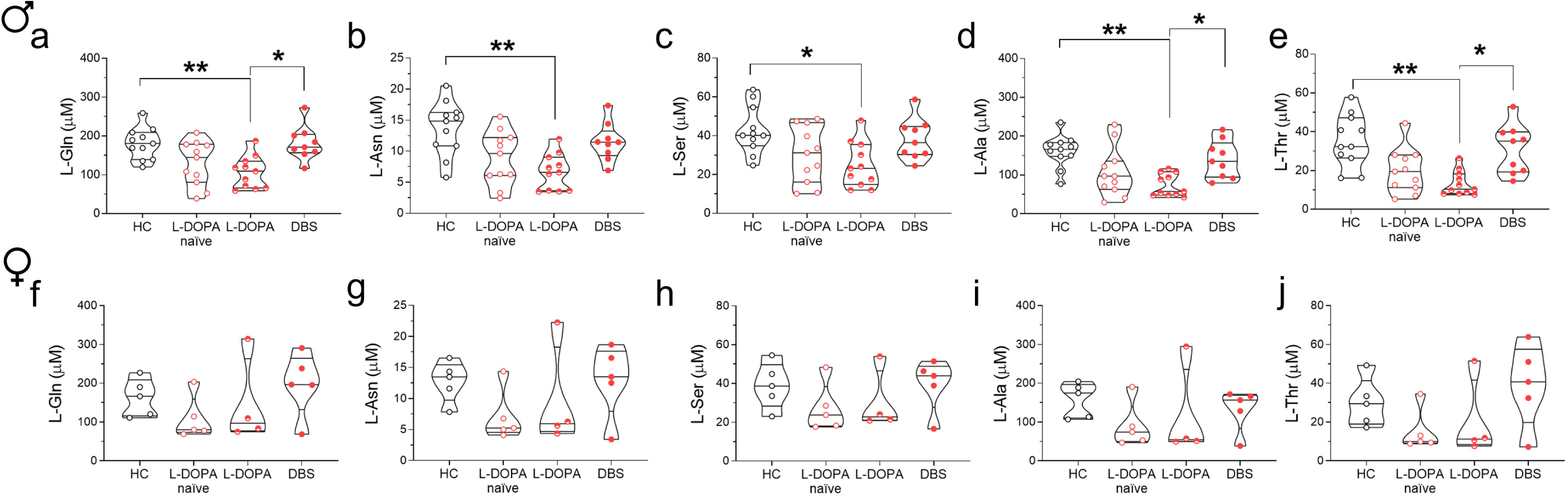
Schematic illustration of circulating neuroactive D- and L-amino acid alterations across PD progression and treatment modalities in male patients. Abbreviations: DBS, Deep Brain Stimulation; PD, Parkinson’s disease; STN, Subthalamic Nucleus.

## Discussion

In our previous metabolomic and HPLC studies in independent cohorts of PD cases and controls, we reported a marked homeostasis disruption in amino acids involved in glutamatergic signalling—particularly NMDAR-related pathways—in the *post-mortem* caudate–putamen, cerebrospinal fluid, and serum of PD patients compared to matched healthy controls (Di Maio et al., 2023; Gervasoni et al., 2025b, 2025a; Imarisio et al., 2024; Nuzzo et al., 2019). Notably, we found that blood amino acid alterations in PD are modulated by sex and genetic background (Yahyavi et al., 2025), with idiopathic male PD patients exhibiting the most pronounced changes relative to both patients carrying established PD pathogenic mutations and female patients (Yahyavi et al., 2025). However, it remains unclear whether these peripheral signatures reflect basal ganglia pathology or originate from systemic metabolic dysfunction. Furthermore, the impact of disease stage and treatment remains undefined, as all participants in our prior studies had comparable disease durations (5–6 years) and were receiving L-DOPA.

Therefore, to address these questions, we quantified circulating glutamatergic system–related neuroactive D- and L-amino acids, along with their metabolic precursors, in male and female healthy controls and PD patients stratified by disease stage and treatment: (1) early-stage L-DOPA-naïve patients treated with MAO-B inhibitors; (2) mid-stage patients treated with L-DOPA; and (3) advanced-stage patients receiving STN-DBS plus L-DOPA.

In agreement with our previous HPLC analyses (Yahyavi et al., 2025), the present study revealed a significant disruption in circulating D- and L-amino acid profiles predominantly in male PD patients, with levels varying markedly according to treatment and disease stage. Although similar directional changes were observed in female PD patients across groups compared to sex-matched healthy controls, these differences did not reach statistical significance—likely due to the limited sample size (N = 4–5). Further validation in larger, sex-stratified cohorts is required to confirm these findings and to clarify the presence and extent of sex-specific differences.

Focusing on male patients, we observed a significant reduction in plasma levels of the NMDAR agonists L-Glu and L-Asp in L-DOPA–naïve PD individuals compared to sex-matched controls. The early reduction of these amino acids is particularly noteworthy, as, in addition to their function as NMDAR agonists, they participate in a broad range of essential cellular biochemical processes. L-Glu is essential for cellular bioenergetics and mitochondrial homeostasis, participates in the urea cycle, contributes to glutathione synthesis, and modulates inflammatory responses (Bjørklund et al., 2021; Du et al., 2016; Yelamanchi et al., 2016). L-Asp is involved in nitrogen disposal, energy metabolism, gluconeogenesis, and nucleotide biosynthesis (Holeček, 2023). Remarkably, the reduction of these neuroactive molecules was detectable within approximately 1.5 years of diagnosis, suggesting that impaired glutamatergic signalling may serve as an early biochemical marker in this subgroup. However, because these patients were receiving MAO-B inhibitors at the time of sampling, further investigation is necessary to disentangle the effects of treatment from disease-driven metabolic changes.

In male patients at mid-stage PD (∼ 7 years’ duration), L-DOPA treatment fails to normalize plasma levels of circulating amino acids. Beyond the reduced L-Asp and L-Glu levels seen in L-DOPA–naïve patients, L-DOPA–treated individuals exhibited further significant decreases in L-Ala, L-Ser, L-Asn and L-Gln compared to sex-matched HC. These findings corroborate our previous report of pronounced amino acid dyshomeostasis in male idiopathic PD patients on L-DOPA treatment (Yahyavi et al., 2025). Additionally, we detected a significant reduction in plasma concentration of L-Thr, an essential amino acid critical for protein synthesis, neurotransmitter production, and immune and gastrointestinal functions (Canfield and Bradshaw, 2019; Mao et al., 2011). While our analysis do not allow as to determine whether these alterations are driven by disease progression, L-DOPA treatment, or their interaction, previous studies have linked changes in circulating amino acid profiles to both disease duration and L-DOPA dose (Figura et al., 2018; LeWitt et al., 2017). Consistent with our observations, these studies have highlighted significant variations in Ala, Arg, Thr, Glu (Figura et al., 2018), and Ser (LeWitt et al., 2017) levels in the serum/plasma of PD patients in relation to disease progression. Notably, Thr and Glu concentrations appear unaffected by L-DOPA dosage (Figura et al., 2018), suggesting that their dysregulation may reflect disease-specific rather than treatment-related mechanisms. Importantly, the functions of D- and L-amino acids examined in the present study extend beyond their roles as modulators of glutamatergic receptors signalling considering they are also critical substrates for the synthesis of proteins and phospholipids, key contributors to mitochondrial bioenergetics, and contribute to antioxidant defence system. Dysfunctions in these processes have been previously implicated in PD pathology (Blesa et al., 2015; Henrich et al., 2023; Watanabe et al., 2025). Taken together, these observations reinforce the notion of PD as a multisystem disorder rather than a disease confined to the central nervous system (Costa et al., 2023).

Lastly, we demonstrate that STN-DBS-therapy, in combination with L-DOPA treatment, can normalize the remarkable dysregulation of circulating neuroactive D- and L-amino acids in patients with advanced PD, characterized by a mean disease duration of approximately 23 years. Notably, in male patients treated with STN-DBS, the plasma concentrations of all amino acids analyzed were comparable to those observed in sex-matched HC. To underscore the magnitude of these effects, we observed that DBS-treated patients exhibited significantly higher plasma levels of D-Ala and Gly than L-DOPA–naïve patients, indicating a reversal of the early-stage biochemical deficits. Although the role of D-Ala in PD remains poorly understood, elevated concentrations of this amino acid have been reported in the grey matter of patients with AD (D’Aniello et al., 1992), and D-Ala supplementation has shown beneficial effects in the treatment of schizophrenia (Tsai et al., 2006). Furthermore, when comparing DBS-treated patients to those receiving L-DOPA, we detected significant higher levels in L-Asp, L-Ala, L-Gln, Gly, and L-Thr, highlighting the capacity of STN-DBS to act on multiple biochemical pathways. Restoration of these neuroactive amino acid levels has important implications for NMDAR signaling, potentially enhancing receptor activation both directly, via L-Asp, and indirectly through the co-agonists D-Ala and Gly (Cummings and Popescu, 2015; Seckler and Lewis, 2020). Importantly, augmented stimulation of the NMDAR glycine-binding site has been associated with significant improvements in motor and non-motor deficits in PD animal models and patients (Frouni et al., 2022, 2021; Gelfin et al., 2012; Schmitz et al., 2013). Moreover, glycine may exert additional effects independent of NMDAR signaling, functioning as a potent inhibitory neurotransmitter via activation of glycine receptors, which are abundantly expressed in key basal ganglia structures, including the caudate-putamen, globus pallidus, and substantia nigra (Waldvogel et al., 2007). Although the underlying mechanisms driving these systemic changes remain to be elucidated, these findings carry two major implications. First, they demonstrate that focal neuromodulation of brain motor circuits via STN-DBS can exert widespread metabolic effects beyond the basal ganglia. Second, they suggest that DBS may confer broad metabolic benefits—beyond its well-established motor symptoms relief—even in patients with long-standing disease. Although, to the best of our knowledge, no study has specifically targeted the impact of STN-DBS on circulating glutamatergic system-related amino acid levels in PD patients stratified for sex, several investigations have reported that STN-DBS can induce significant systemic effects in individuals with advanced PD. Notably, this intervention has been associated with marked increases in body weight and BMI within the first year post-surgery (Barichella et al., 2003; Macia et al., 2004; Novakova et al., 2007; Węgrzynek-Gallina et al., 2025)—a feature also observed in our STN-DBS-treated subgroup of advanced PD patients. While the mechanisms underlying this weight gain remain debated, emerging evidence suggests that STN-DBS may influence systemic lipid and glucose metabolism (Batisse-Lignier et al., 2013; Carrillo et al., 2024; Węgrzynek-Gallina et al., 2025), supporting the hypothesis of a functional crosstalk between the central nervous system and peripheral metabolic regulation.

Notably, recently in the study of Carrillo et al. (2024), authors analyzed plasma lipid and metabolite profiles in the same PD and healthy control cohort examined in the present study, identifying the 50 most discriminant lipid species between cases and controls (Carrillo et al., 2024). These lipids were markedly upregulated in both L-DOPA–naïve and L-DOPA–treated patients relative to controls, whereas STN-DBS–treated patients exhibited lipid profiles closely resembling those of healthy individuals. In conjunction with our data, these results imply that STN-DBS in advanced PD patients may exert systemic metabolic benefits by normalizing both lipidomic and amino acid perturbations. However, Carrillo et al. also reported amino acid alterations that differ from our HPLC findings (Carrillo et al., 2024). This apparent discrepancy likely reflects differences in cohort design (pooled versus sex-stratified) and analytical methodology (HPLC versus gas chromatography– time-of-flight MS).

We acknowledge that the present study has limitations, including the relatively small cohort size, absence of longitudinal follow-up, and lack of stratification based on the genetic background of PD patients, which may contribute to heterogeneity in metabolic profiles. Additionally, the PD subgroups differ not only in treatment modality but also in disease stages.

In summary, our study delineates a progressive disruption of circulating neuroactive amino acids in PD evident from the earliest disease stages and exacerbating over disease progression. Notably, L-DOPA treatment fails to correct these biochemical imbalances, whereas STN-DBS in advanced PD patients normalizes D- and L-amino acid levels to those of sex-matched HC. Collectively, these findings highlight the potential of circulating amino acid dysregulation as an early biomarker of PD and reveal the systemic metabolic benefits of STN-DBS beyond its motor symptom–alleviating effects.

## Materials and Methods

### PD cohort population

804 independent and unrelated PD patients (501 males; 300 familiar and 504 sporadic cases) were recruited at the IRCCS Mediterranean Neurological Institute (MNI) in Pozzilli (Carrillo et al., 2024; Gialluisi et al., 2021, 2019; Palomba et al., 2023; Tirozzi et al., 2021). All PD subjects were of European ancestry and were evaluated by neurologists of the Parkinson Study Group, according to published diagnostic criteria (Postuma et al., 2015). The PD cohort was recruited from June 2015 to December 2017, and from June 2021 to December 2022, with a thorough protocol comprising neurological examination and evaluation of non-motor domains. Information about family history, demographic characteristics, anamnesis, and pharmacological therapy was also collected. The Movement Disorder Society revised version of the Unified Parkinson’s Disease Rating Scale Part III (33 items, maximum score 132; hereafter called UPDRS) was used to assess clinical motor symptoms (Goetz et al., 2008). These included language, facial expressions, tremor, rigidity, agility in movements, stability, gait and bradykinesia. Cognitive abilities were tested through an Italian validated version of the Montreal Cognitive Assessment (MoCA) (Conti et al., 2015). Cognitive domains assessed include short-term memory (5 points); visuospatial abilities via clock drawing (3 points), and a cube copy task (1 point); executive functioning via an adaptation of Trail Making Test Part B (1 point), phonemic fluency (1 point), and verbal abstraction (2 points); attention, concentration, and working memory via target detection (1 point), serial subtraction (2 points), digits forward and backward (1 point each); language via confrontation naming with low-familiarity animals (3 points), and repetition of complex sentences (2 points); and orientation to time and place (6 points). The total score was given by the sum of these domains and then divided by the maximum score that could be obtained (30 points). Non-motor symptoms were assessed through an Italian validated version of Non Motor Symptoms Scale (NMS) for PD (Chaudhuri et al., 2006). This scale tests 9 items, including cardiovascular domain, sleep/fatigue, mood/cognition, perceptual problems/hallucinations, attention/memory, gastrointestinal, urinary, sexual function, and ability to taste or smell. Among the PD patients, we used the 14 patients from the cohort who had had a DBS implant (10 patients with a bilateral implant and 4 with a monolateral implant) for a comparable time at the time of recruitment. Patients carrying GBA1 mutations were excluded. To compare the data of an equal number of patients per group, we selected 16 patients in the initial phase of the disease who were not yet being treated with L-DOPA and 15 patients who had been being treated with L-DOPA for at least 5 years. Written informed consent was obtained from all participants. The ethical board of the IRCCS Neuromed approved the study protocols: N°9/2015, N°19/2020 and N°4/2023.

### Collection and storage of serum samples

Blood sampling was performed after a 6-h fasting. Blood samples from PD patients and healthy subjects were collected in blood collection tubes (BD Vacutainer, K2E (EDTA) then, after two centrifugations at 1900g for 10 min, plasma was aliquoted and stored at − 80 °C. Unique anonymized codes have been assigned to the samples for processing and subsequent analysis, maintaining the confidentiality of personal data.

### HPLC analysis of amino acids content

Plasma samples (100 μl) were mixed in a 1:10 dilution with HPLC-grade methanol (900 μl) and centrifuged at 13,000 ×g for 10 min. Supernatants were dried and then suspended in 0.2 M trichloroacetic acid (TCA). TCA supernatants were then neutralized with 0.2 M NaOH and subjected to precolumn derivatization with o-phthaldialdehyde/N-acetyl-L-cysteine in 50% methanol. Amino acids derivatives were resolved on a UHPLC Nexera X3 system (Shimadzu) by using a Shimpack GIST C18 3-μm reversed-phase column (Shimadzu, 4.0 × 150 mm) under isocratic conditions (0.1 M sodium acetate buffer, pH 6.2, 1% tetrahydrofuran, and 1 ml/min flow rate). A washing step in 0.1 M sodium acetate buffer, 3% tetrahydrofuran and 47% acetonitrile, was performed after every run. Identification and quantification of amino acids were based on retention times and peak areas, compared with those associated with external standards. The detected amino acids concentration was expressed as μM.

### Statistical analysis

Clinical and demographic characteristics were described using, as summary statistics, median and the interquartile range (IQR) or absolute frequencies. Comparisons between PD groups and HC were evaluated using Kruskal-Wallis’ test for demographic and clinical parameters. Comparison of plasma amino acid levels was performed using Kruskal-Wallis’ test followed by adjusted Dunn’s post hoc multiple comparisons, when required. ANCOVA model with age and LEDD as covariate was used to control for the effect of these factors on plasma amino acid levels. Significance was set at p < 0.05 for all analyses.

## Conflict of Interest

The authors declare that the research was conducted in the absence of any commercial or financial relationships that could be considered as a potential conflict of interest.

## Data availability

All data are available in the main text.

## Author contributions

TN: Investigation and data analysis; FC: Data analysis; MS: Writing – review & editing; CG: Investigation; ADM: Investigation; AP: Investigation; SP: Review & editing; NM; review & editing; FE: review & editing; TE: Funding acquisition, Project administration, Supervision– review & editing. AU: Conceptualization, Funding acquisition, Project administration, Supervision, Writing – review & editing.

## Acknowledgments

This study was partially funded by Italian Ministry of University and Research (PRIN 2022 - COD. 2022XF7YYL_02 to AU and PRIN 2022 – COD. 2022W3RKLJ to TE). The work of A.U., T.N. and T.E. was supported by NEXTGENERATIONEU (NGEU) and funded by the Ministry of University and Research (MUR), National Recovery and Resilience Plan (NRRP), project MNESYS (PE0000006) – A Multiscale integrated approach to the study of the nervous system in health and disease (DN. 1553 11.10.2022).

The work of T.E. was supported by Next Generation EU - PNRR M6C2 Investimento 2.1 valorizzazione e potenziamento della ricerca biomedica del SSN grant n. PNRR-MAD-2022-12375960 and grant n. PNRR-MCNT2-2023-12377375. TE was also supported by Ministry of Health, Ricerca Corrente.

The authors are grateful to all the patients, their caregivers, the Clinical Parkinson’s Disease Center of IRCCS Pozzilli and the PD biobank of IRCCS Neuromed and IGB-CNR for the kind cooperation with this study.

## Abbreviations

CSF: cerebrospinal fluid
D-Ala: D-alanine
D-Ser: D-serine
DBS: deep brain stimulation
Gly: glycine
HC: healthy controls
HPLC: high-performance liquid chromatography
L-Ala: L-alanine
L-Asn: L-asparagine
L-Asp: L-aspartate
L-Gln: L-glutamine
L-Glu: L-glutamate
L-Thr: L-threonine
LEDD: levodopa equivalent daily dose
LID: L-DOPA-induced dyskinesia
MPTP: 1-methyl-4-phenyl-1,2,3,6-tetrahydropyridine
NMDAR: N-methyl-D-aspartate receptor
NMS: non-motor symptoms
PD: Parkinson’s disease
SNpc: substantia nigra pars compacta
STN: subthalamic nucleus
UPLC-MS: ultra-performance liquid chromatography–mass spectrometry.

## Notes

### Competing Interest Statement

The authors have declared no competing interest.

### Summary of Updates

This version of the manuscript has been revised to improve the writing style and correct typos

